# Cryo-EM structures of chronic traumatic encephalopathy tau filaments with PET ligand flortaucipir

**DOI:** 10.1101/2022.12.15.520545

**Authors:** Yang Shi, Bernardino Ghetti, Michel Goedert, Sjors H.W. Scheres

## Abstract

Positron emission tomography (PET) imaging allows monitoring the progression of amyloid aggregation in the living brain. [^18^F]-Flortaucipir is the only approved PET tracer compound for the visualisation of tau aggregation. Here, we describe cryo-EM experiments on tau filaments in the presence and absence of flortaucipir. We used tau filaments isolated from the brain of an individual with Alzheimer’s disease (AD), and from the brain of an individual with primary age-related tauopathy (PART) with a co-pathology of chronic traumatic encephalopathy (CTE). Unexpectedly, we were unable to visualise additional cryo-EM density for flortaucipir for AD paired helical or straight filaments (PHFs or SFs), but we did observe density for flortaucipir binding to CTE Type I filaments from the case with PART. In the latter, flortaucipir binds in a 1:1 molecular stoichiometry with tau, adjacent to lysine 353 and aspartate 358. By adopting a tilted geometry with respect to the helical axis, the 4.7 Å distance between neighbouring tau monomers is reconciled with the 3.5 Å distance consistent with π-π-stacking between neighbouring molecules of flortaucipir.

## Introduction

Accumulation of assembled tau protein is the hallmark of multiple neurodegenerative diseases that are collectively known as tauopathies [1]. Specific clinical and neuropathological features are used to define and distinguish the tauopathies, that include Alzheimer’s disease (AD), primary age-related tauopathy (PART), chronic traumatic encephalopathy (CTE), progressive supranuclear palsy (PSP), corticobasal degeneration (CBD) and Pick’s disease (PiD). Electron cryo-microscopy (cryo-EM) imaging has allowed atomic structure determination of tau filaments from *post mortem* brain of patients in recent years [2–7] Distinct conformers of tau (or tau folds) define different diseases and provide a structure-based classification of tauopathies [7]. In adult human brain, six tau isoforms are expressed [8]. Three tau isoforms have 3 microtubule-binding repeats (3R), whereas the other three have 4 microtubule-binding repeats (4R). In several tauopathies, all six (3R+4R) tau isoforms assemble into filaments. The Alzheimer and CTE folds are the only known tau folds for 3R+4R tauopathies. Depending on their inter-protofilament interfaces, two identical protofilaments with the Alzheimer fold assemble as paired helical filaments (PHFs) or as straight filaments (SFs) in AD and PART [2, 4, 9]. Likewise, two identical protofilaments assemble as type I and type II filaments in CTE [5].

The ability to detect tau pathology in the living brain is essential for understanding the relationship between neuropathology and clinical symptoms and for monitoring the effects of mechanism-based therapies. Several tau tracer molecules have been developed for visualizing the spatiotemporal distribution of filamentous tau deposits in the brains of living subjects using positron emission tomography (PET) imaging. The degree and patterns of tau PET retention strongly overlap with regions affected by brain atrophy [10, 11] and correlate with concurrent cognitive performance [12]. Tau PET has shown excellent diagnostic [13–16] and prognostic [17] performance and potential in longitudinal studies [18–20]. [^18^F]-Flortaucipir [21] (also known as [^18^F]-T807, [^18^F]-AV-1451, or TAUVID) is the most widely used tau PET tracer in the clinic and it was the first (in May 2020) and is so far the only compound to be approved for this purpose by the U.S. Food and Drug Administration.

Flortaucipir discriminates between AD and other tauopathies [13, 22]. Patterns of [^18^F] flortaucipir retention *in vivo* reflect *post mortem* Braak staging, in support of PET-based staging of AD [23]. Increased retention of [^18^F]-flortaucipir was also observed in non-AD cases, such as CTE [24–27], CBD [28], and PSP [29, 30], although its usefulness as a biomarker in these diseases is less clear. Despite studies using autoradiography [31, 32], binding assays [33] and molecular dynamics simulations [34, 35], it remains unclear how flortaucipir binds to tau filaments. This lack of experimental structural information impedes structure-based development of better and more disease-specific PET ligands.

Using cryo-EM, we previously identified two binding sites for another tau PET tracer compound, APN-1607 or PM-PBB3 (propanol modification of pyridinyl-butadienyl-benzothiazole 3), in the β-helix of PHFs and SFs, and a third site in the C-shaped cavity of SFs [9]. Here, we used similar methods to examine the binding of flortaucipir to Alzheimer and CTE tau filaments, from a case of AD and a case of PART with additional CTE pathology.

## Results

### Cryo-EM structures of tau filaments with flortaucipir from a case of AD

We first performed cryo-EM structure determination of tau filaments from the sarkosyl-insoluble fractions of the frontal cortex from a case with AD (case 2 in reference [4]). Prior to cryo-EM grid preparation, we incubated sarkosyl-insoluble fractions with flortaucipir at 3 nmol per gram of brain tissue to saturate potential binding to the tau filaments (+flortaucipir). As controls, we imaged tau filaments that were incubated with DMSO without flortaucipir (-flortaucipir). Using helical reconstruction in RELION [36, 37], we determined the structures of PHFs to resolutions of 2.7 Å for both the +flortaucipir and -flortaucipir structures. Comparison of both maps (Figure 1) did not identify additional densities in the +flortaucipir map that were separated from those of tau, and the corresponding difference map did not show peaks at a threshold of 5 standard deviations. Due to their low abundance (less than 5% of the tau filaments), we were unable to obtain useful reconstructions of SFs.

**Figure 1.**
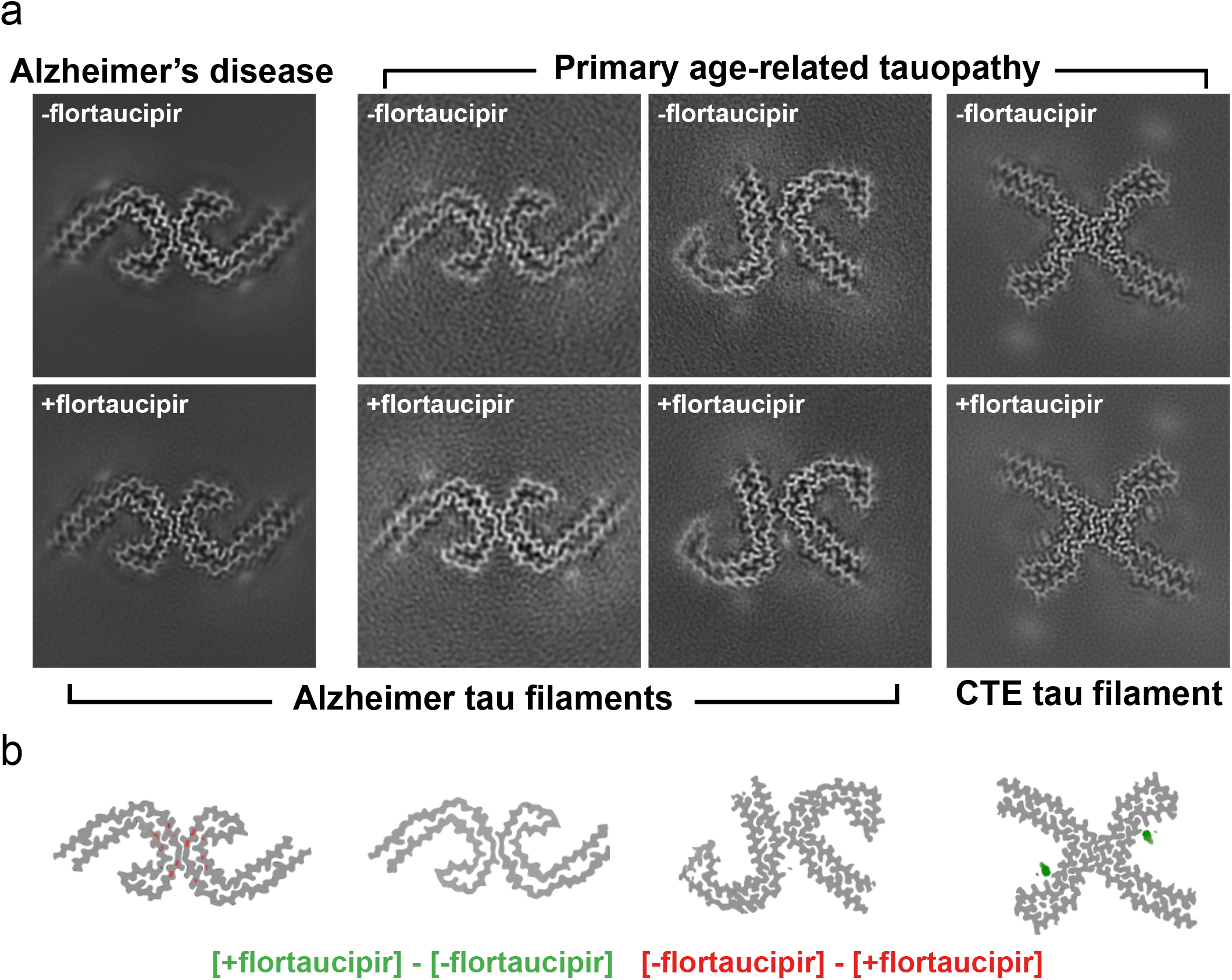
Cryo-EM structures of tau filaments with flortaucipir. a. XY-cross-sections, with a projected depth of approximately one β-rung, for cryo-EM reconstructions of tau filaments with and without flortaucipir (+flortaucipir and -flortaucipir, respectively) for PHFs from the case with AD (first column from the left), for PHFs from the case with PART (second column); for SFs from the case with PART (third column); and CTE Type I filaments from the case with PART (fourth column). b. Corresponding difference maps ([+flortaucipir] — [-flortaucipir]) for the same four columns; positive difference density is shown in green at a threshold of 5 standard deviations in the difference map; negative difference density is shown in red at a threshold of 5 standard deviations.

### Cryo-EM structures of tau filaments with flortaucipir from a case of PART with a copathology of CTE

To exclude the possibility that flortaucipir only binds to Alzheimer SFs, we also performed structure determination for tau filaments from the hippocampus of a case of PART (case 1 in reference [9]) that was previously shown to have more SFs than PHFs [9]. To increase the accessibility of flortaucipir to the ordered core of tau filaments, we pronase-treated the sarkosyl-insoluble fractions before incubation with flortaucipir at 10 nmol per gram of brain tissue to saturate potential binding to the tau filaments (+flortaucipir). We determined structures of SFs to 2.6 Å (+flortaucipir) and 2.7 Å (-flortaucipir) and of PHFs to 3.4 Å (+flortaucipir) and 3.7 Å (–flortaucipir). Again, we did not identify additional densities in the maps with flortaucipir, for PHFs or SFs, and the corresponding difference maps did not show any peaks at a threshold of 5 standard deviations (Figure 1).

For the case of PART, which had a co-pathology of CTE, we also observed CTE type I filaments, for which we determined structures to 2.6 Å (+flortaucipir) and 2.7 Å (-flortaucipir). When comparing these maps, we did identify an additional density in the C-shaped cavity adjacent to K353 and D358. The additional density extended up 13 standard deviations in the corresponding difference map. When displayed at a threshold of 5 standard deviations (Figure 1), the difference density was well separated along the 4.8 Å rungs of the β-sheets along the filaments, with an angle of approximately 46 degrees between the long axis of the additional density and the helical axis. The molecular structure of flortaucipir fits well into the elongated shape of the additional density (Figure 2). The intensity of the additional density is comparable to the density of the filament, suggesting a close to 1:1 molecular stoichiometry between flortaucipir and tau. The flat appearance of the additional density suggests that the nonhydrogen atoms in each flortaucipir molecule adopt a planar configuration that is characteristic of a π-conjugated system. The 46-degree orientation between the planar structure of the flortaucipir molecules and the helical axis of the tau filaments leads to an interplanar distance of 3.5 Å, which is consistent with π-π-stacking between adjacent flortaucipir molecules. The interaction between flortaucipir and tau is probably driven by hydrogen bonding between the nitrogen in the six-membered benzene ring of the gamma-carboline moiety and the hydroxyl of D358 with an N-O distance of 2.9 Å. In this orientation, the flortaucipir molecule also fits the additional density better than with the same nitrogen pointing towards K353 (Supplementary Figure 2). The overall structure of the tau filaments in the +flortaucipir and -flortaucipir maps is identical, indicating that the binding of flortaucipir does not induce conformational changes within the CTE filaments upon binding (Figure 3a).

**Figure 2.**
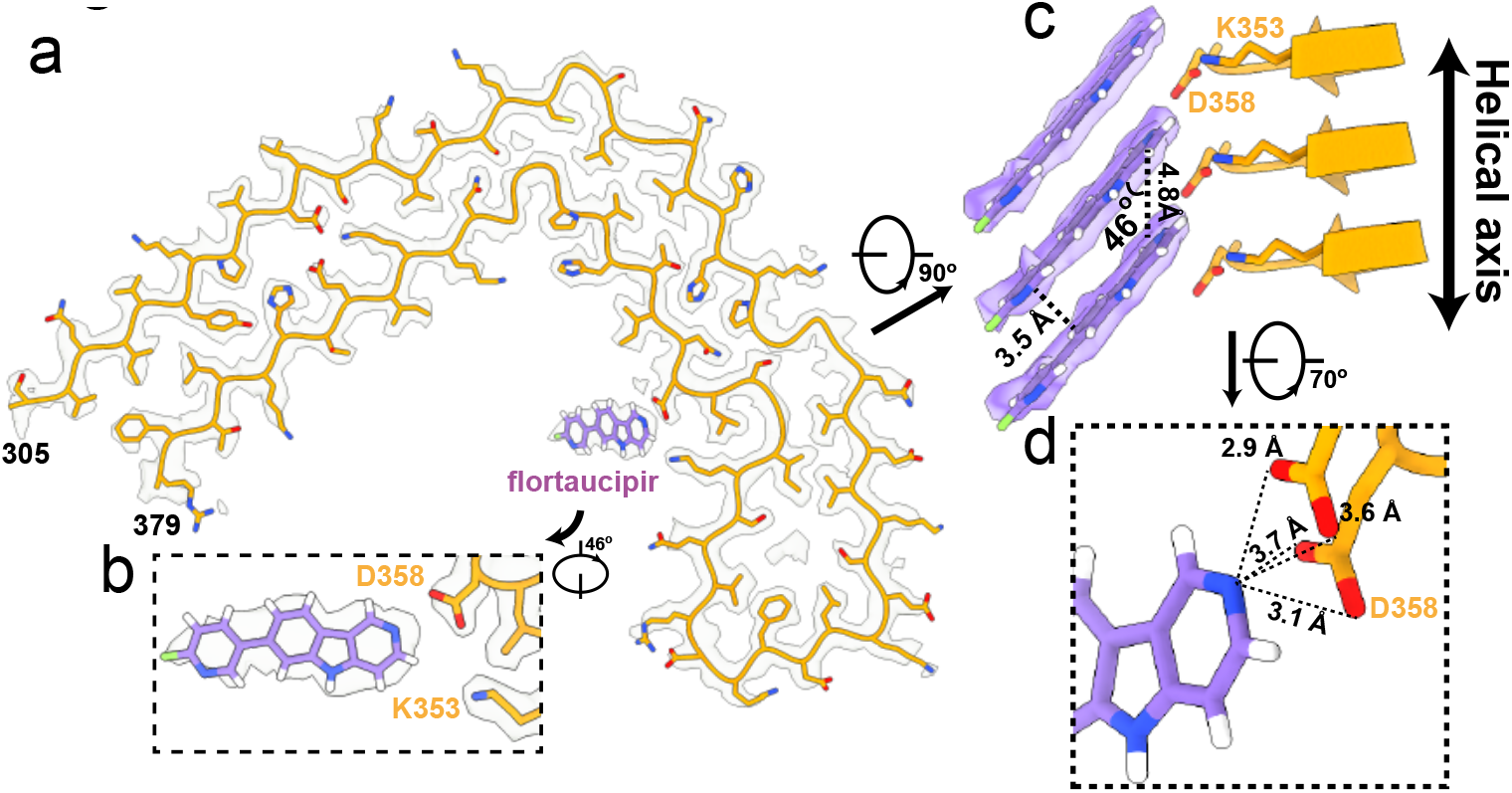
Flortaucipir binding site to CTE Type I filaments. a. Cryo-EM density (white) with the atomic model for tau (orange) and flortaucipir for the CTE Type I filaments from the case with PART. b. Rotated, zoomed-in view of the flortaucipir binding site. c. Side view of the flortaucipir binding site. d. Zoomed-in view to highlight the distances between the nitrogen atom in the six-membered benzene ring of the gamma-carboline moiety and the oxygen atoms of D358.

**Figure 3.**
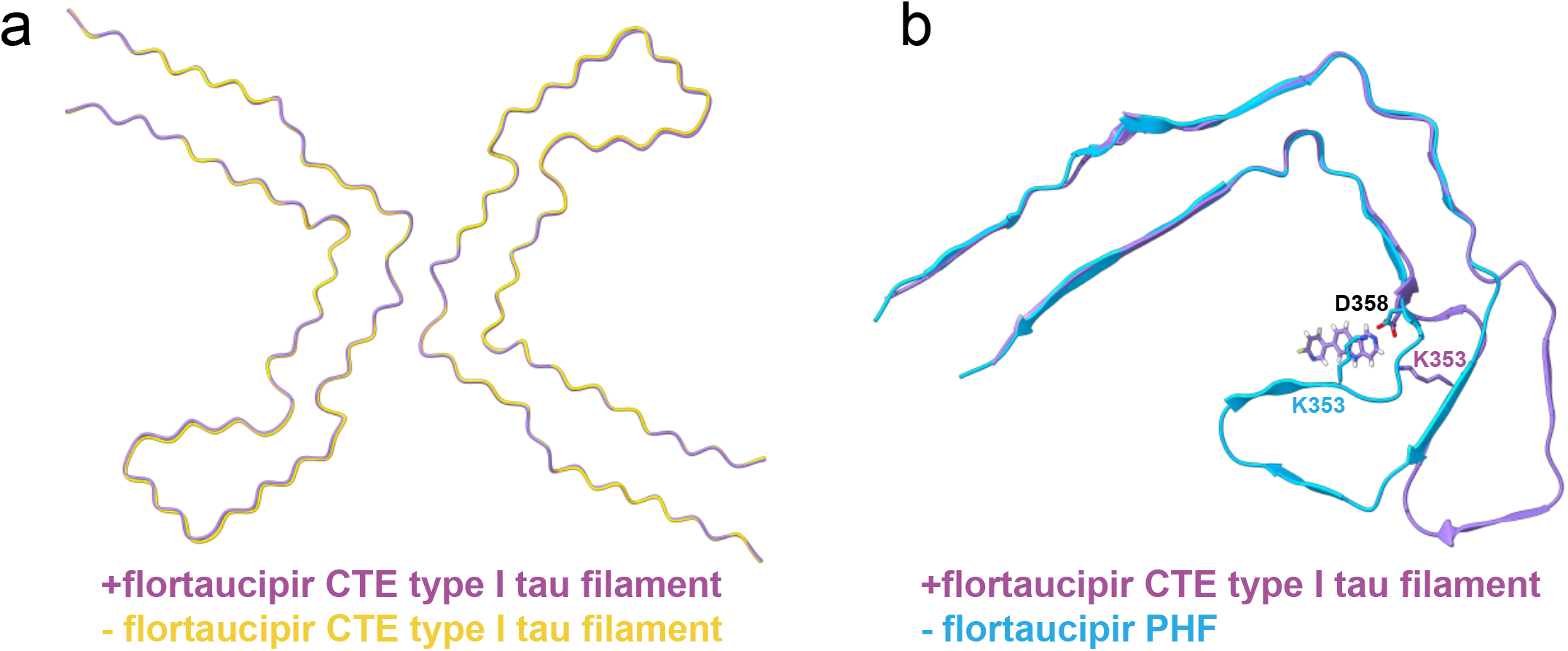
Structure comparisons. a. Comparison of the structures of the CTE Type I filaments from the case with PART, with and without flortaucipir (+flortaucipir and -flortaucipir, respectively), indicate that the tau filaments do not change upon flortaucipir binding. b. Comparison of the structures of the AD and CTE Type 1 filaments from the case with PART with flortaucipir suggests that the more closed C-shape structure of the AD fold cannot accommodate flortaucipir in the same way at the CTE Type I filament.

## Discussion

Our cryo-EM results are in apparent contradiction with observations from the clinical use of [^18^F] flortaucipir, as it has been approved for AD. Fortaucipir has not yet been approved for CTE. In one study, autoradiography revealed binding to *post mortem* tissue sections of multiple cases of CTE, but with varying degrees of specificity, including off-target binding to monoamine oxidase-A [38]. Using similar experiments, another study reported faint or no binding of flortaucipir that could be attributed to tau aggregates [39]. In a study of eleven cases with traumatic encephalopathy syndrome, tau-PET imaging with flortaucipir showed either mildly elevated or no frontotemporal binding, and mildly elevated medial temporal binding in a subset of cases, with values being considerably lower than in AD [40]. Tau-PET imaging of 26 former National Football League players with cognitive and neuropsychiatric symptoms showed that tau standardized uptake value ratios were higher than for 31 controls, but without association between tau deposition and scores on cognitive and neuropsychiatric tests [26]. The clinical usefulness of flortaucipir for CTE thus remains unclear.

Yet, when comparing the cryo-EM maps of tau PHFs from a case of AD incubated with flortaucipir with those incubated without flortaucipir, we did not observe significant differences. The same was also true for PHFs and SFs from a case of PART. However, CTE type I filaments from the same case of PART did show a strong (up to 13 standard deviations above the noise) additional density, with a shape and size that fits well with the binding of a single molecule of flortaucipir to each molecule of tau in the CTE type I filaments.

How can these observations be reconciled with what is known about the clinical usefulness of flortaucipir? Why did cryo-EM not detect binding of flortaucipir to PHFs and SFs, when it is useful for tau-PET imaging in AD? One possible explanation is that flortaucipir binds to the disordered fuzzy coat of the tau filaments, where it would be invisible for cryo-EM reconstruction. Although we cannot exclude this explanation, we do not favour it for two reasons. First, it is unclear how flortaucipir could bind in a specific manner to intrinsically disordered parts of the tau protein. Secondly, most cases of end-stage AD have predominantly extracellular tangles [1], which lack most of the fuzzy coat, and which would thus not be detected by flortaucipir. It has been reported that flortaucipir labels extracellular tangles in AD brains [33]. A second explanation, which we prefer, is that flortaucipir binds to AD tau filaments in highly sub-stoichiometric amounts. Cryo-EM reconstruction relies on the averaging over many tau molecules that make up the filaments. The result is a reconstruction in which the density for every rung of tau molecules in the amyloid filament has been forced to be identical. (In PHFs, additional symmetry between the two protofilaments is also imposed so that all tau molecules have identical density.) Therefore, sub-stoichiometric binding of flortaucipir would lead to a decrease in its reconstructed density, until it gets drowned in the noise.

Although we observed stoichiometric binding of flortaucipir to CTE type I filaments, flortaucipir tau-PET imaging results in CTE have so far not been conclusive. It is possible that the accessibility for flortaucipir of tau filaments in brain may be less in CTE than in AD. We also note that we examined flortaucipir binding to tau filaments from an individual with PART who had a CTE co-pathology, rather than from an individual with CTE.

Despite these uncertainties, our results may further our understanding of the binding of small-molecule compounds to amyloid filaments. The more open C-shaped cavity of the CTE fold compared to the Alzheimer fold may explain why PHFs and SFs did not bind flortaucipir in the same manner (Figure 3b). Only two other cryo-EM studies of amyloid filaments in complex with small-molecule compounds have been published so far. We previously reported cryo-EM reconstructions of tau-PET ligand PM-PBB3 to PHFs and SFs [9], whereas Seidler et al [41] reported the structure of PHFs in complex with epigallocatechin gallate (EGCG), a compound that disaggregates PHFs. Each of these studies revealed different binding modes of the small molecule compounds. PM-PPB3, a ~20 Å long molecule with an extended π system, was found to bind to multiple sites on PHFs and SFs, with individual molecules binding parallel to the helical axis spanning up to six rungs of tau molecules. EGCG, a molecule with a benzenediol ring adjoined to a tetrahydropyran moiety, a galloyl ring and a pyrogallol ring, binds with 1:1 stoichiometry to PHFs, at the interface between protofilaments. Interestingly, as the relative angle between the main aromatic rings of EGCG and the helical axis is closer to 90 degrees, the interplanar distance between consecutive EGCG molecules is larger than the optimal distance for π-π-stacking, which was hypothesized to contribute to its disaggregating properties. The 46-degree angle observed between flortaucipir and the helical axis of CTE type I filaments may be a hallmark of amyloid-binding compounds that form intermolecular π-π-stacking interactions. In agreement with these observations, a similar stacking arrangement (with a 44-degree angle) was reported on bioRxiv for the binding of PET tracer GTP-1 to AD PHFs, while we were preparing this manuscript [42].

In conclusion, we describe the mode of binding of flortaucipir to CTE type I filaments from a case of PART with CTE co-pathology. Unexpectedly, we could not detect binding of flortaucipir to PHFs and SFs from the same case of PART and a case of AD.

## Acknowledgements

We are grateful to Eli Lilly for providing flortaucipir. We thank Drs Kelly Conway and Suchira Bose for helpful discussions; the EM facility at the Medical Research Council (MRC) Laboratory of Molecular Biology for help with cryo-EM data acquisition; and Toby Darling and Jake Grimmett for help with high-performance computing. This work was supported by the UK Medical Research Council (MC_UP_A025_1013 to S.H.W.S. and MC_U105184291 to M.G.) and the US National Institutes of Health (NIA AG P30 010133 to B.G.). For the purpose of open access, the MRC Laboratory of Molecular Biology has applied a CC BY public copyright licence to any Author Accepted Manuscript arising.

## Additional Information

For the purpose of open access, the authors have applied a CC-BY public copyright license to any Author Accepted Manuscript version arising.

## Materials and Methods

### Extraction of tau filaments

Sarkosyl-insoluble material was extracted from the frontal cortex of an individual with AD (case 2 in reference [4]) and from the hippocampus of an individual with PART (case 1 in reference [9]). For the AD case, tissues were homogenized in 10 vol (w/v) extraction buffer consisting of 10 mM Tris—HCl, pH 7.5, 10% sucrose, 0.8 M NaCl, 5 mM EDTA, 1 mM EGTA and a protease and phosphatase inhibitor (Thermo Fisher). Homogenates were spun at 20,000 g for 20 min and supernatants were retained. Pellets were homogenized in 5 volumes (w/v) extraction buffer and centrifuged at 20,000g for 20 min. Both supernatants were combined, brought to 1% sarkosyl and incubated for 60 min at room temperature. Following a 60 min centrifugation at 100,000g, pellets were resuspended in 250 μl/g extraction buffer and spun at 20,000g for 20 min. The resulting supernatants were centrifuged at 100,000 g for 1 h. For the PART case, tissues were homogenized in 20 vol (w/v) extraction buffer. Homogenates were brought to 2% sarkosyl and incubated for 30 min at 37 °C. Following a 10 min centrifugation at 10,000g, the supernatants were spun at 100,000g for 60 min. The pellets were resuspended in 700 μl/g extraction buffer and centrifuged at 10,000g for 10 min. The supernatants were diluted threefold in 50 mM Tris-HCl, pH 7.4, containing 0.15 M NaCl, 10% sucrose and 0.2% sarkosyl, and spun at 100,000g for 60 min. For cryo-EM, sarkosyl-insoluble pellets were resuspended in 30 and 100 μl/g of 20 mM Tris-HCl, pH 7.4 containing 100 mM NaCl, for the AD and PART extracts respectively.

### Binding of flortaucipir to tau filaments

A 10 mM solution of flortaucipir (provided by Eli Lilly) was prepared in anhydrous dimethyl sulfoxide (DMSO, Thermo Fisher) and stored at −20 °C. Sarkosyl-insoluble brain material was incubated with 100 μM flortaucipir in 20 mM Tris—HCl, pH 7.5, 100 mM NaCl, 2% DMSO, for 3 h at room temperature. Controls were incubated with buffer containing 2% DMSO.

### Electron cryo-microscopy

Samples were applied to glow-discharged holey carbon gold grids (Quantifoil R1.2/1.3, 300 mesh), and plunge frozen in liquid ethane using an FEI Vitrobot Mark IV. Images of the samples from the AD case used Gatan K2 summit and K3 detectors in counting mode on a Titan Krios (Thermo Fisher) at 300 kV for +flortaucipir and -flortaucipir samples respectively. A GIF quantum energy filter (Gatan) was used with a slit width of 20 eV to remove inelastically scattered electrons. Images of the samples from the PART case were acquired using a Falcon-4 detector without energy filter in counting mode on a Titan Krios (Thermo Fisher) at 300 kV. Further details are given in Supplementary Table 1. To increase the accessibility of flortaucipir to the core of tau filaments, samples (with and without flortaucipir) of the case with PART were treated with 0.1 mg/ml of pronase for 30-40 min, prior to making grids [43].

### Helical reconstruction

Movie frames were gain-corrected, aligned, dose-weighted and then summed into a single micrograph using RELION’s own motion correction program [44]. Aligned and non-dose weighted micrographs were used to estimate the contrast transfer function (CTF) by CTFFIND-4.1 [45]. All subsequent image-processing steps were performed using helical reconstruction methods in RELION [36, 37]. Tau filaments were picked manually, and extracted using an inter-box distance of 14.1 Å. For reference-free 2D classification, segments with a box size comprising an entire helical crossover were downscaled by a factor of 3 to speed up calculations. Different types of filaments were separated by reference-free 2D classifications and segments contributing to suboptimal 2D averages discarded. Initial 3D models were constructed *de novo* from 2D class averages comprising an entire helical crossover using the relion_helix_inimodel2d program [37]. Helical twists were estimated by crossover distances from 2D class averages. Segments for 3D auto-refinement were then re-extracted using a box containing about 45% helical crossover, without downscaling. With an initial 3D model that was low-pass filtered to 10 Å, 3D auto-refinement was carried out for several rounds with optimization of helical twist and rise, after reconstructions had shown separation of β-strands along the helical axis. We then performed CTF refinement, followed by 3D classification without further image alignment, to remove segments that yielded suboptimal 3D reconstructions. Final reconstructions were sharpened using standard postprocessing procedures in RELION [44]. Overall resolution estimates were calculated from the Fourier shell correlations (FCSs) at 0.143 between two independently refined half-maps, using phase-randomisation to correct for convolution effects of a generous, soft-edged solvent mask that extended to 20% of the height of the box. Using the relion_helix_toolbox program [37], helical symmetries were imposed on the post-processed maps. For further details, see Supplementary Figure 1 and Supplementary Table 1.

### Model building and refinement

Protein Data Bank (PDB) entry 6NWP [5] was used as initial reference for atomic model building of CTE type I filaments from the case of PART. Models containing three β-sheet rungs were refined in real-space by PHENIX [46] with local symmetry. MolProbity [47] was used for model validation. To confirm the absence of overfitting, FSC curves between one half-map and the model, which was refined against the other half-map, were checked. Additional details are given in Supplementary Figure 1 and Supplementary Table 1. The final map and model of CTE type I filaments from the case of PART with flortaucipir were submitted to the Electron Microscopy Data Bank (accession number EMDB-16329) and the Protein Data Bank (accession number 8BYN), respectively.

## Figure Legends

**Supplementary figure 1.**
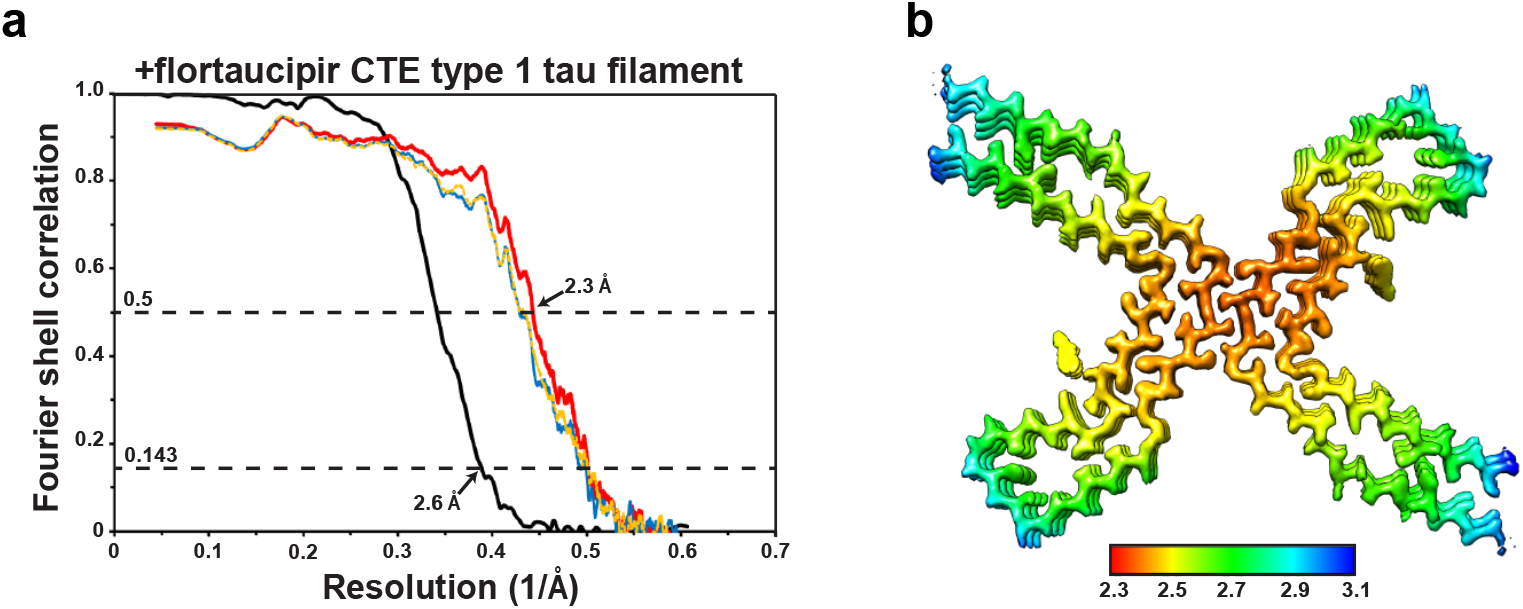
Cryo-EM and model resolution assessment. a. Fourier shell correlation (FSC) curves for cryo-EM maps of the CTE Type I filaments with flortaucipir (in black); for the final refined atomic model against the final cryo-EM map (in red); for the atomic model refined in the first half map against that half map (in blue); and for the refined atomic model in the first half map against the other half map (in yellow). B. Top view of the cryo-EM density coloured by local resolution (in Å).

**Supplementary figure 2.**
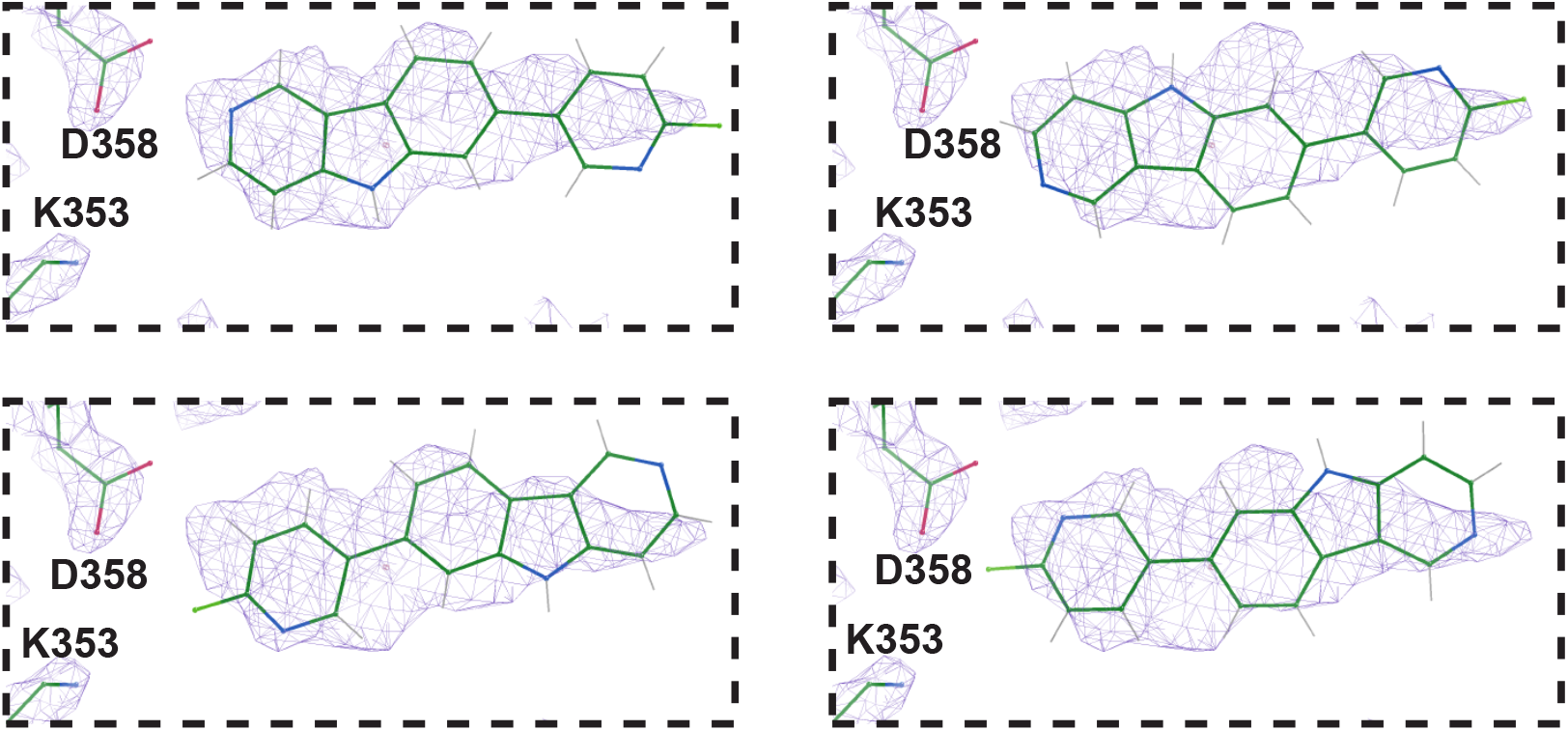
Alternative dockings of flortaucipir. Four alternatives for the docking of flortaucipir into the additional cryo-EM density are shown. We favour the top left conformation, which positions the nitrogen atom in the six-membered benzene ring of the gamma-carboline moiety in close proximity of the oxygen atoms of D358 and provides the best fit to the density.

**Supplementary Table 1.** Cryo-EM data collection, refinement and validation statistics

|  | AD<br>(+flortaucipir) | AD<br>(-flortaucipir) | PART<br>(+flortaucipir) | PART<br>(-flortaucipir) |
| --- | --- | --- | --- | --- |
| **Data collection and processing** |  |  |  |  |
| EM type | Titan Krios | Titan Krios | Titan Krios | Titan Krios |
| Magnification | 105,000 | 96,000 | 96,000 | 96,000 |
| Detector | K2 | K3 | Falcon 4 | Falcon 4 |
| Voltage (kV) | 300 | 300 | 300 | 300 |
| Electron exposure (e-/Å<sup>2</sup>) | 60 | 51 | 40 | 40 |
| Defocus range (μm) | -1.0 to -2.5 | -1.8 to -3.0 | -1.0 to -2.0 | -1.0 to -2.0 |
| Pixel size (Å) | 1.15 | 0.828 | 0.824 | 0.824 |
| Symmetry imposed | C1 | C1 | C1 | C1 |
| Initial particle images (no.) | 673,528 | 656,157 | 385,082 | 1,123,147 |
| Final particle images (no.) | 114,174 | 561,329 | PHF:4,792<br>SF:42,917<br>CTE1: 58,019 | PHF:2,207<br>SF:36,686<br>CTE1: 48,163 |
| Map resolution (Å) | 2.70 | 2.68 | PHF:3.43 | PHF:3.66 |
| FSC threshold(0.143) |  |  | SF: 2.62<br>CTE1: 2.60 | SF: 2.70<br>CTE1: 2.69 |
| Helical twist (°) | 179.43 | 179.43 | PHF:179.46<br>SF: -1.08<br>CTE1: 179.43 | PHF:179.47<br>SF: -1.08<br>CTE1: 179.42 |
| Helical rise (Å) | 2.37 | 2.47 | PHF:2.37<br>SF: 4.77<br>CTE1: 2.37 | PHF:2.38<br>SF: 4.77<br>CTE1: 2.37 |
| **Refinement** |  |  |  |  |
| Initial model used (PDB code) |  |  | 6NWP |  |
| Model resolution (Å) |  |  | 2.3 |  |
| FSC threshold (0.5) |  |  |  |  |
| Map sharpening *B* factor (Å<sup>2</sup>) |  |  | -70 |  |
| Model composition |  |  |  |  |
| Non-hydrogen atoms |  |  | 3564 |  |
| Protein residues |  |  | 450 |  |
| Ligands |  |  | 6 |  |
| *B* factors (Å<sup>2</sup>) |  |  |  |  |
| Protein |  |  | 20.23 |  |
| Ligand |  |  | 12.17 |  |
| R.m.s. deviations |  |  |  |  |
| Bond lengths (Å) |  |  | 0.006 |  |
| Bond angles (°) |  |  | 0.680 |  |
| Validation |  |  |  |  |
| MolProbity score |  |  | 1.35 |  |
| Clashscore |  |  | 6.29 |  |
| Poor rotamers (%) |  |  | 0.00 |  |
| Ramachandran plot |  |  |  |  |
| Favored (%) |  |  | 98.63 |  |
| Allowed (%) |  |  | 1.37 |  |
| Disallowed (%) |  |  | 0.00 |  |

